# T cells use TNF and IFNγ for paracrine killing with target discrimination programmed by pathogen-derived factors

**DOI:** 10.64898/2025.12.23.696308

**Authors:** Matthew C. Guttormson, Erica N. Krogman, Allison R. Carr, Skylar A. Rizzo, Josey Muske, Maia Ziaee, Nicholas R. Therrien, Nicholas D. Sun, Romina Ghale, Michael J. Medlyn, Daniel D. Billadeau, Larry R. Pease, Kathryn A. Knoop, Adrian T. Ting

## Abstract

Cytotoxic T lymphocytes (CTLs) are known to eliminate target cells through perforin-mediated, contact-dependent killing - a process limited by the number of effector T cells despite its serial nature. CTLs likely also possess a mass killing mechanism that can eliminate target cells more efficiently. Using CAR-T cells against B7H3 present on B16 tumor targets, we demonstrate that activated T cells can also kill targets in a perforin-independent manner by releasing diffusible TNF and IFNγ capable of killing nearby targets in a paracrine fashion even if they bear no antigen. Against an unperturbed target, paracrine killing is inefficient but is enhanced by the deletion of TNFR1 signaling molecules such as TAK1, HOIP, or TBK1/IKKε. Notably, these molecules are inhibited naturally by pathogen-encoded antagonists. Expression of a microbial-encoded antagonist, such as the *Yersinia*-encoded YopJ that antagonizes TAK1 or Ebola-encoded VP35 that antagonizes TBK1/IKKε, alters the target cell response to these cytokines from non-lethal to lethal. We propose that target cells are inherently resistant to killing by TNF and IFNγ, but the presence of a microbial factor alters this sensitivity, providing for the selective elimination of the infected cell while minimizing harm to uninfected bystander cells. We propose the term ‘pathogen-restriction’ to describe this discriminatory mechanism. Potentially, any microbial-derived factor that crosstalks with the TNF and IFNγ signaling pathways can function as a pathogen-restriction element.

**GRAPHICAL ABSTRACT:** 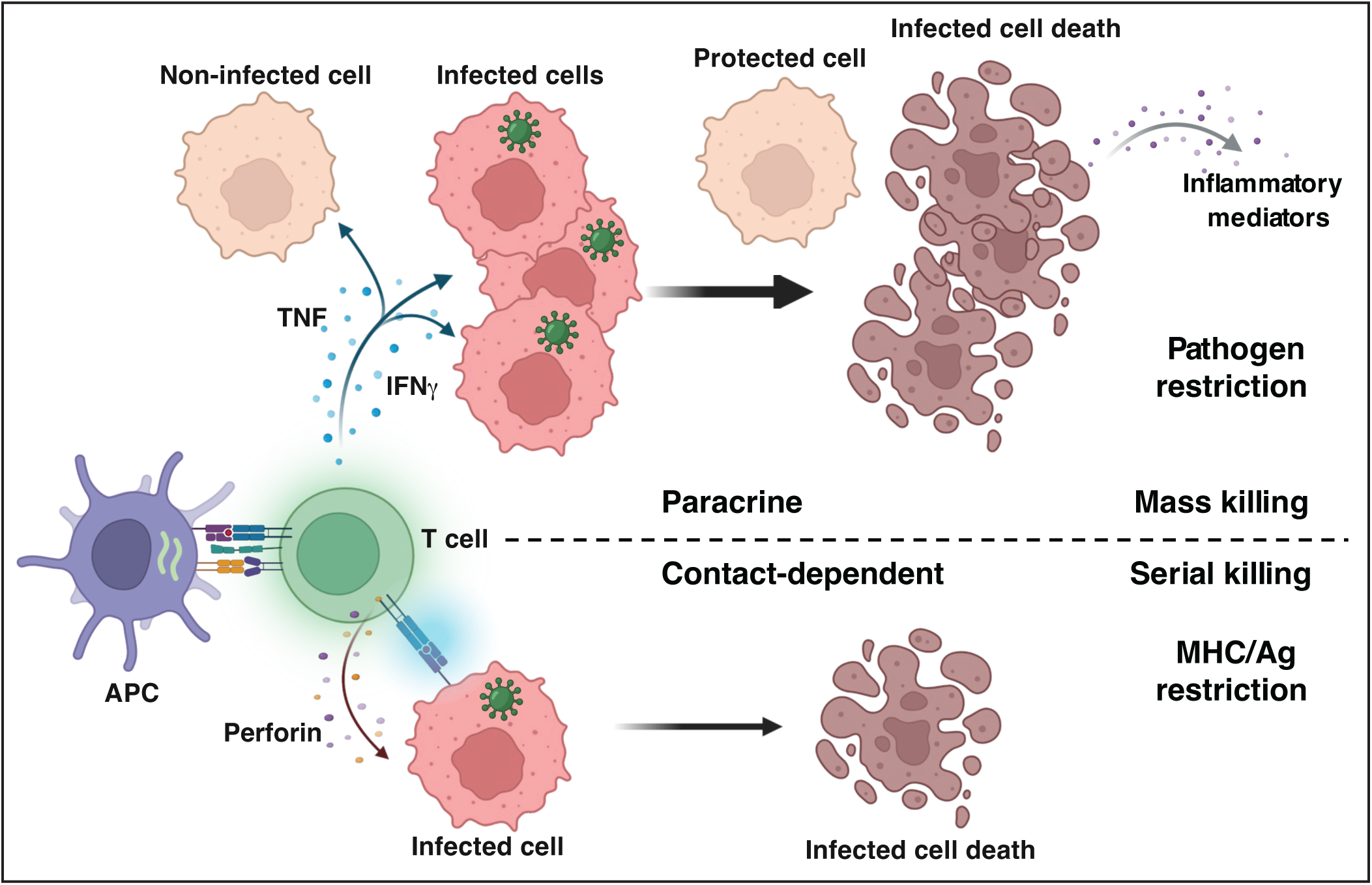

**Short Summary:** T cells use TNF and IFNγ to kill target cells in a paracrine manner where target elimination is restricted by the presence of pathogen-derived factors.

## INTRODUCTION

T cells are central to adaptive immunity, orchestrating responses that eliminate infected or malignant cells through antigen recognition and cytotoxic effector functions (*1, 2*). Conventional models emphasize perforin/granzyme-mediated killing at the immunological synapse—a highly directional process requiring intimate contact between cytotoxic T lymphocytes (CTLs) and their targets (*3–5*). This serial killing paradigm imposes inherent limitations: each CTL can only kill one target at a time, and imaging studies have shown this to be inefficient (*6*). With the frequency of naïve T cells encoding a TCR specific for a microbial-derived antigen estimated to be low (*7*), coupled with a rapidly replicating pathogen, contact-dependent killing is likely insufficient. While this can be mitigated by clonal expansion, our immune system has very likely evolved a more efficient method of eliminating infected cells where the interaction of a single T cell with a single antigen-bearing cell can be leveraged to eliminate many more infected cells. A guiding principle in such a mass elimination mechanism must remain that of specificity in which only the infected cells, but not uninfected bystanders, are removed. A conceptual framework that allows us to understand how this leveraging is achieved during an anti-microbial response will have a significant clinical impact. This is especially important in cancer immunotherapy where contact-dependent killing mediated by perforin/granzyme is likely also insufficient for control against proliferating tumor cells by the limited number of tumor-specific T cells. Here, using CAR-T cells and B16 F1 murine melanoma cells as a model target, we demonstrate that T cells deploy a paracrine killing mechanism capable of mass elimination of target cells in a discriminatory but antigen-agnostic manner. We propose that discriminatory killing is programmed by the presence of pathogen-derived factors in the intended target and suggest the term ‘pathogen-restriction’ to describe this discriminatory mechanism.

## RESULTS

### T cells can kill targets via TNF and IFNγ independent of perforin

The initial question we asked was whether T cells can kill target cells without perforin. We utilized CAR-T cells because we can generate them from wild-type (WT) or perforin-deficient *Prf1^-/-^* (PRF1 KO) splenocytes (Fig. S1A). The CAR was based on the m276 scFv specific for B7H3/CD276 (*8*), an extracellular protein expressed on many tumor cells (*9*), including B16 F1 melanoma cells that we used as a model target (Fig. S1B). When co-cultured with B16 cells at an E:T ratio of 1:1 over an extended period of 3 days, WT CAR-T cells against B7H3 mediated robust target cell killing (Fig. 1A & 1B). PRF1 KO CAR-T cells retained similar killing capacity, inducing cell death under identical conditions albeit with slower kinetics (Fig. 1A & 1B), demonstrating the existence of a perforin-independent killing mechanism in T cells. We confirmed this with PRF1 KO CAR-T cells against hCD19 based on the FMC63 scFv using B16 cells transduced with hCD19 as targets (Fig. S1C). Since activated T cells express TNF and IFNγ, and the two cytokines can synergize to kill tumor cells and non-tumor cells (*10, 11*), we tested if these two cytokines are involved in the perforin-independent killing of B16 targets. Anti-TNF and anti-IFNγ had no effect on killing by WT CAR-T cells but completely abrogated killing by PRF1 KO CAR-T cells (Fig. 1A & 1B), confirming the role of the two cytokines in the alternate perforin-independent killing. High concentration of recombinant TNF (100 ng/mL) by itself, but not IFNγ, was able to induce discernable death of B16 targets, but combination of the two cytokines had a synergistic effect (Fig. 1C & 1D). Genetic ablation of FADD or STAT1 in B16 cells conferred resistance to killing by cytokine treatment (Fig. 1E) and by PRF1 KO CAR-T cells (Fig. 1F), implicating TNFR1-mediated death-signaling and IFNγR-mediated transcription in cytokine-induced death. Biochemical analysis indicated that TNF and IFNγ synergized to induce apoptosis evidenced by cleaved CASP8 and CASP3, as well as pyroptosis evidenced by cleaved GSDMD (Fig. 1G). Blotting of lysates generated from B16 targets co-cultured with B7H3 CAR-T cells also showed the appearance of cleaved CASP8 p18 fragment and cleaved GSDMD with both WT and PRF1 KO T cells but reversed by anti-TNF and anti-IFNγ (Fig. 1H). Cytokine blockade did not inhibit the appearance of cleaved CASP3 and PARP in co-cultures from WT CAR-T cells, consistent with those targets undergoing perforin/granzyme-mediated apoptosis. Both cleaved CASP8 p18 and GSDMD in response to cytokines or CAR-T cells were diminished in B16 targets with FADD and STAT1 ablation (Fig. S1D & S1E). These results indicate that high levels of TNF and IFNγ or ratio of T cells can kill B16 tumor targets.

**Fig. 1.**
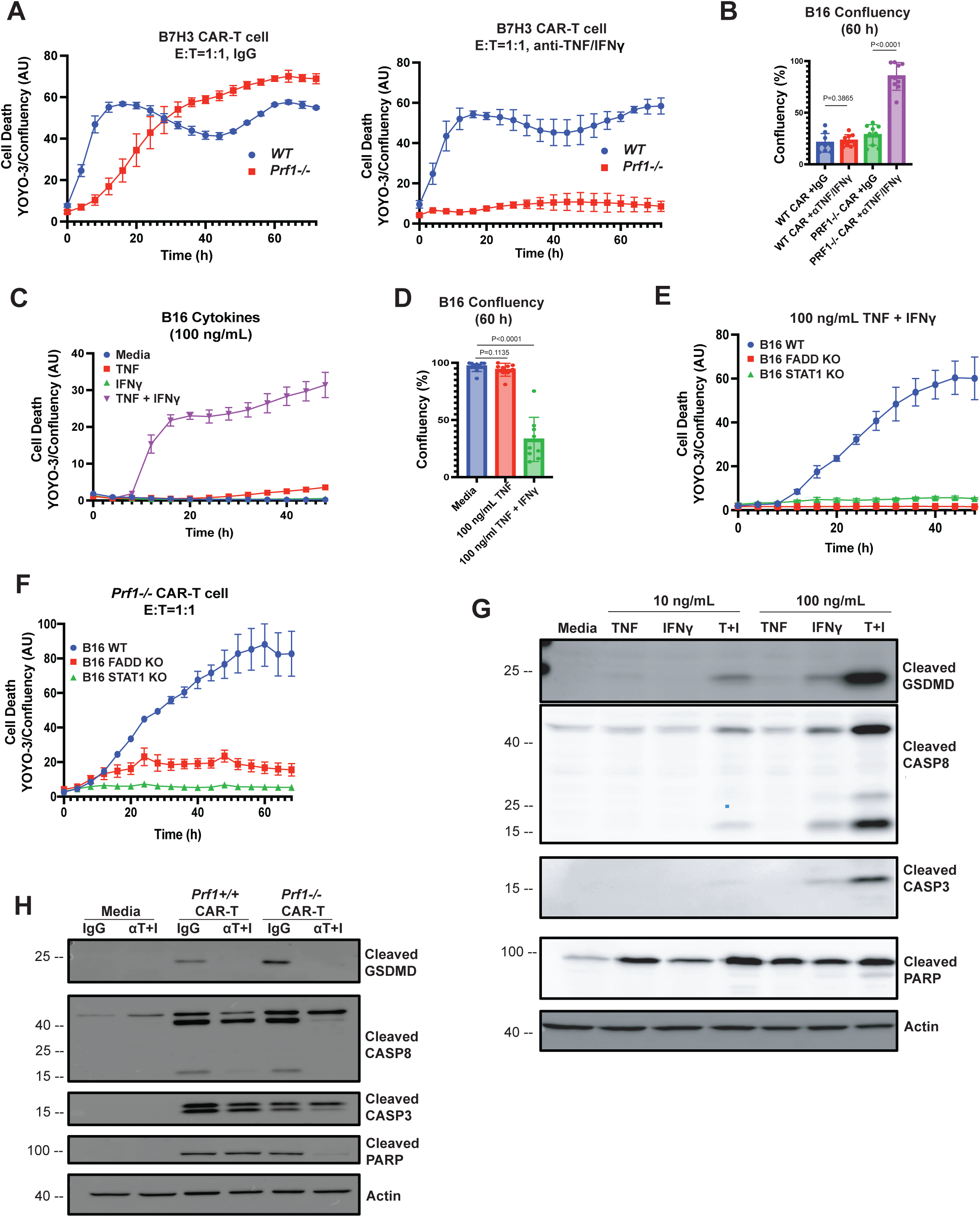
T cells kill targets via TNF and IFNγ independent of perforin. **(A)** WT and *Prf1-/-* anti-B7H3 CAR-T cells were co-cultured with B16 melanoma cells in a 96W plate in triplicates. An E:T ratio of 1:1 was used. Control IgG or neutralizing anti-TNF and anti-IFNγ (50 ug/mL) were added to the wells. Target viability was assessed by YOYO-3 staining and live-cell imaging for 72 h using the IncuCyte S3 instrument. Confluency of the cells in each well were also quantified. Data is presented as YOYO-3 counts normalized to confluency in each well in arbitrary units. Values are triplicate mean ± SD. Data from one representative experiment is shown. **(B)** The experiment in (A) was independently performed 3 times. The mean confluency at 60 h of 9 wells from the 3 biological replicates is shown in the bar chart. (**C)** B16 cells were stimulated in triplicates with 100 ng/mL TNF, IFNγ, or both, and imaged in the IncuCyte for 48 h in the presence of YOYO-3. Values are triplicate mean ± SD. Data from one representative experiment is shown. **(D)** The experiment in (C) was performed 3 times. The mean confluency of 9 wells at 60 h from the 3 biological replicates is shown in the bar chart. (**E)** WT, FADD KO or STAT1 KO B16 cells were treated with a combination of 100 ng/mL TNF and IFNγ. Cell death was analyzed for 48h in the IncuCyte as in (A). Representative data from one experiment is shown. **(F)** WT, FADD KO or STAT1 KO B16 cells were co-cultured with *Prf1*-/- CAR-T cells against B7H3 at an E:T ratio of 1:1. Target killing was analyzed as in (A). (**G**) Western blot analysis of the indicated proteins in B16 cells stimulated for 24 h with TNF and IFNγ as indicated. **(H)** B16 cells were co-cultured with *Prf1-/-* CAR-T cells at 1:1 ratio in the presence of control IgG or antibodies against TNF and IFNγ (50 ug/mL). Data in (G, H) are representative of 3 independent experiments. *p* values from Mann-Whitney tests are shown for data in (B, D).

### T cells use cytokines for inflammatory paracrine killing

The existence of an alternate mode of killing suggests that it performs a function distinct from perforin-mediated killing. Unlike perforin-mediated killing, which is rapid, the slower kinetics of target death induced by TNF and IFNγ (Fig. 1A), and the fact that TNF and IFNγ activate NFκB- and STAT1-dependent transcription that can synergize to induce inflammatory gene transcription (*12, 13*), suggest that these dying cells may emit inflammatory cues. Blotting of nuclear extracts from B16 cells demonstrated that simultaneous nuclear localization of NFκB and STAT1 transcription factors occurred in cells treated with a combination of both cytokines (Fig. 2A). We then performed gene expression analysis with the NanoString nCounter platform using the host response panel, which showed a synergistic effect of the two cytokines on a subset of genes (Fig. S2), including the inflammatory chemokines CCL2 and CXCL9 (Fig. 2B). ELISA of culture supernatants from B16 cells treated with TNF and IFNγ confirmed CCL2 and CXCL9 production (Fig. 2C). While pyroptosis might provide some level of inflammatory cues, the pronounced inflammatory gene induction by combined TNF and IFNγ suggests that these dying targets will exert an effect on downstream responses.

**Fig. 2.**
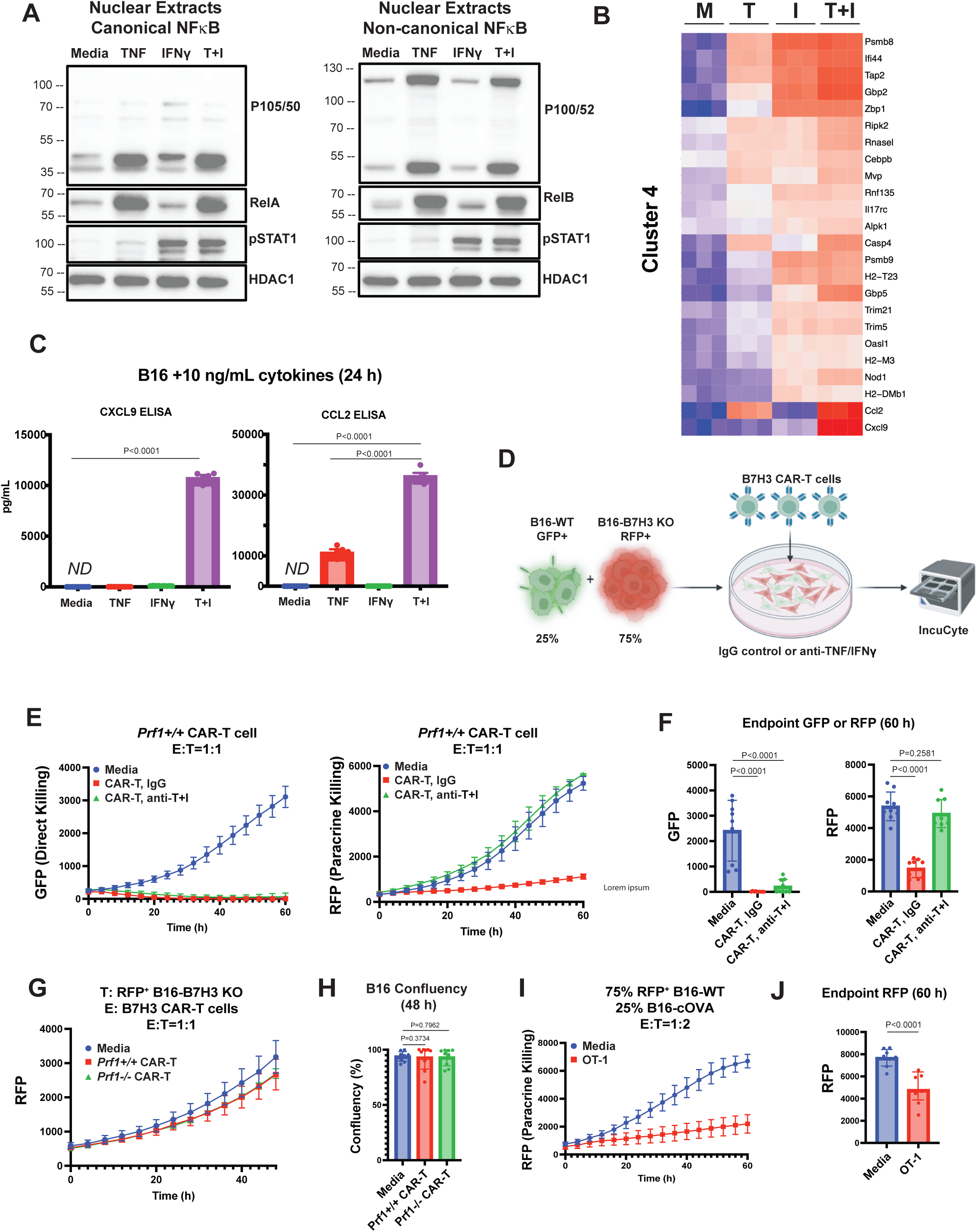
Cytokine-mediated paracrine killing by T cells is inflammatory. **(A)** Nuclear extracts from B16 cells treated for 24 h with 10 ng/mL TNF, IFNγ, or both were blotted for canonical and non-canonical NFκB and STAT1. HDAC1 was used as a loading control. **(B)** Biological triplicate samples of RNA isolated from B16 cells treated as in (A) were analyzed using the Nanostring nCounter Host Response panel. Heatmap shows a group of genes from unsupervised clustering whose expression was synergistically induced by TNF and IFNγ. **(C)** Supernatants from B16 cells treated as in (A) were analyzed for level of CCL2 and CXCL9 by ELISA. Values are the mean ± SD from three independent experiments, each performed in technical triplicates. **(D)** Schematic of paracrine killing assay. B16 WT cells were stably transduced with nuclear green fluorescent protein (GFP) and B16 B7H3 KO cells with nuclear raspberry fluorescent protein (RFP). The two population of targets were mixed and co-cultured with CAR-T cells against B7H3. **(E)** Mixed GFP⁺ WT and RFP⁺ B7H3 KO B16 cells (1:3 ratio) were co-cultured with *Prf1+/+* CAR-T cells at E:T ratio of 1:1. Control IgG or anti-TNF/IFNγ (50 ug/mL) was also added. GFP and RFP counts in each well was quantified by IncuCyte live-cell imaging for 60 h. Values are mean ± SD from triplicate well. Data from one representative experiment is shown. **(F)** Experiment in (E) was performed three times. Bar graphs represent mean GFP and RFP counts in the wells at endpoint from the three biological replicates. **(G)** RFP+ B16 B7H3 KO cells were co-cultured with media, *Prf1+/+* or *Prf1-/-* CAR-T cells against B7H3 at an E:T of 1:1 and imaged for 48 h. **(H)** Mean confluency of wells at endpoint from three independent experiments in (G). **(I)** 1:3 mixture of B16-cOVA and RFP⁺ B16 WT cells were co-cultured with OT-1 T cells at an E:T ratio of 1:2. RFP counts in each well was quantified by IncuCyte imaging for 60 h. Values are triplicate mean ± SD. **(J)** Experiment in (I) was performed three times. Bar graphs represent mean RFP counts in the wells at endpoint from three biological replicates. *p* values from Mann-Whitney tests are shown for data in (C, F, H and J).

Since TNF and IFNγ are small diffusible proteins, another property distinct from perforin-containing supramolecular attack particles (SMAPs) (*14*), T cells could potentially use this mechanism to kill a target in a paracrine fashion, without directly engaging with it. We analyzed paracrine killing by mixing GFP-labeled WT B16 cells with raspberry fluorescent protein (RFP)-labeled B7H3 KO B16 target cells at a 1:3 ratio (Fig. 2D) and co-cultured them with B7H3 CAR-T cells at varying E:T ratios. Live-cell imaging in the IncuCyte showed that WT CAR-T cells eliminated both the GFP+ and RFP+ target populations, demonstrating their ability to kill target cells that do not express the CAR antigen (Fig. 2E & 2F). Addition of anti-TNF and anti-IFNγ reversed the elimination of the antigen-negative RFP-labeled cells but not the GFP-labeled B7H3-expressing targets (Fig. 2E & 2F). Note that cultures containing only RFP-labeled B7H3-negative targets were not killed by CAR-T cells (Fig. 2G & 2H) indicating that the presence of antigen-expressing cells is required to first activate the CAR-T cells in order to eliminate the antigen-negative targets. To confirm that paracrine killing is not an artefact of CAR-T cells, we mixed non-labeled B16 cells expressing cytoplasmic OVA (cOVA) with RFP-labeled B16 cells that do not express cOVA and co-cultured these mixed targets with OT-1 T cells. Like CAR-T cells, OT-1 T cells were also able to kill antigen-negative B16 targets (Fig. 2I & 2J). Thus, distinct from perforin-mediated killing, T cells use TNF and IFNγ to kill targets in an inflammatory paracrine manner, including targets that do not express the cognate antigen.

### Target discrimination during paracrine killing is conferred by altering TNFR1 death signaling

Since TNF and IFNγ can bind to any cell and could potentially trigger its demise, this is likely to be dangerous. Our immune system must have evolved a mechanism to selectively eliminate an infected cell but leave uninfected bystander cells unharmed. We noted that in our CAR-T cell co-culture killing assays, we used relatively high E:T ratios, which are unlikely to be attained during an anti-microbial or anti-tumor response. We also had to use high concentrations of the recombinant cytokines to trigger cell death of WT B16 targets. Therefore, we postulate that the selectivity issue can be solved if in the default (uninfected) state, TNF and IFNγ are non-lethal to a cell, but in a cell that harbors a pathogen, the presence of a microbial-derived factor alters their signaling and converts the response to a lethal one. Several pathogens encode molecules that antagonize TNFR1 signaling molecules as part of their strategy to block induction of NFκB/MAPK/IRF3-regulated inflammatory genes. These include YopJ from Yersinia pestis that blocks TAK1 (*15*), VP35 from Ebola virus that inhibits TBK1/IKKε (*16*), and IpaH1.4/2.5 from Shigella flexneri that blocks HOIP (*17*). Deficiency in TAK1, TBK1/IKKε and HOIP are known to switch the TNF response from survival to RIPK1-mediated death (*18–22*). We hypothesize that the presence of these microbial-encoded factors marks a cell for killing by TNF and IFNγ. We tested this by deleting TAK1, HOIP, or TBK1/IKKε in B16 targets to mimic their blockade by pathogen-encoded molecules (Fig. S3A). These knockouts (KO) markedly increased sensitivity to combined TNF and IFNγ compared to WT targets (Fig. S3B & S3C). Under limiting conditions at lower E:T ratios, PRF1 KO CAR-T cells were unable to kill WT targets but effectively killed the three KO targets (Fig. 3A & 3B). The three KO targets displayed different sensitivity to CAR-T cell killing, with TAK1 and TBK1/IKKε deficiencies having a more potent effect (Fig. 3A & 3B). Antibody blockade confirms killing of the KO targets is mediated by TNF and IFNγ (Fig. 3C). Blockade of individual cytokines suggest that IFNγ had a more potent role, as its neutralization had a greater effect on reversing killing as compared to neutralizing TNF (Fig. S3D). Biochemical analysis with the TBK1/IKKε DKO and TAK1 KO targets indicate that they were sensitized to T cell killing via apoptosis (Fig. S3E).

**Fig. 3.**
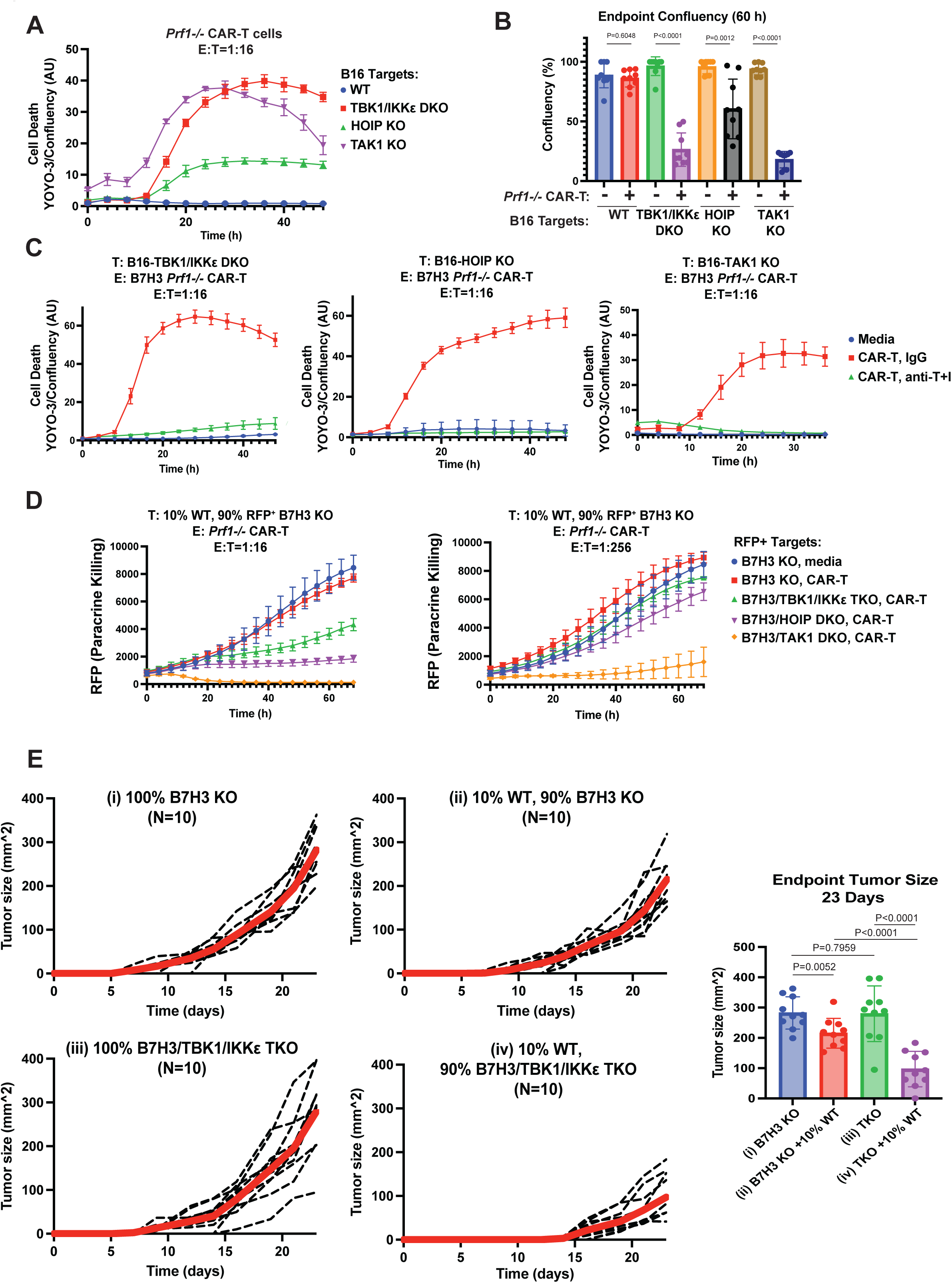
Target discrimination during paracrine killing is conferred by modulation of TNFR1 signaling. **(A)** *Prf1-/-* CAR-T cells were co-cultured at varying E:T ratios with B16 targets that are WT, TAK1 KO, TBK1/IKKε DKO and HOIP KO, and analyzed in the IncuCyte for 48 h. Cell death data for E:T of 1:16 is shown. **(B)** The experiment in (A) was independently performed 3 times. The mean confluency at 60 h of 9 wells from the 3 biological replicates is shown in the bar chart. **(C)** The indicated B16 targets were co-cultured with media or with *Prf1-/-* CAR-T cells at varying E:T ratios with control IgG or anti-TNF/IFNγ (50 ug/mL). Target killing was analyzed in the IncuCyte for 48 h and data for E:T of 1:16 is shown. **(D)** RFP-labeled B7H3 KO targets of the indicated genotypes were mixed with 10% unlabeled B7H3⁺ B16 WT cells and co-cultured with *Prf1*-/- CAR-T cells at varying E:T ratios. Paracrine killing was quantified by RFP counts of the B7H3-negative targets in the IncuCyte. Data from E:T ratios of 1:16 and 1:256 are shown. **(E)** 2.5 x 10^5^ B7H3 KO or B7H3/TBK1/IKKε TKO B16 cells that were unmixed or mixed with 10% WT B16 cells were implanted into B6 hosts. 5 days later, mice were administered *Prf1-/-* CAR-T cells against B7H3 via tail vein infusion and tumor growth measured. Spider plots shown were compiled from two independent experiments, each with N=5 per group. Thick red line indicates average tumor volume in each group of mice. Bar chart shows mean tumor volume ± SD for each group at day 23. *p* values from Mann-Whitney tests are shown for data in (B and E).

The above experiments do not directly assess paracrine killing as all targets express B7H3 so they can engage with the CAR-T cells. To do so, we ablated B7H3 from the three KO targets and their WT counterparts (Fig. S4A) and labeled them with RFP. The RFP-labeled targets were then ‘spiked’ with 10% unlabeled B7H3-expressing WT B16 cells, and co-cultured with *Prf1^-/-^* CAR-T cells against B7H3. Despite the absence of the B7H3 antigen, these knockout targets but not their WT counterpart were effectively killed under limiting T cell condition (Fig. 3D). This effect was particularly pronounced in TAK1 knockout cells, which were efficiently killed at E:T ratios as low as 1:256 (Fig. 3D). Paracrine killing was completely abrogated following TNF and IFNγ blockade (Fig S4B), confirming its dependence on cytokines. Titrating the amount of antigen-expressing B16 cells spiked into cultures of antigen-deficient targets showed that even a frequency of 2.5% is sufficient to trigger paracrine killing of antigen-negative TAK1- and TBK1/IKKε-deficient targets (Fig. S4C). Cell-free supernatants from CAR-T cells co-cultured with B7H3-positive, but not B7H3-negative targets, also preferentially kill KO targets (Fig. S4D-F) providing further support for paracrine killing. To confirm that paracrine killing is a general feature of T-cell cytotoxicity rather than a CAR-specific phenomenon, we spiked RFP-labeled WT or TBK1/IKKε DKO B16 targets with 10% unlabeled OVA-expressing B16 and co-cultured the mixture with *Prf1^-/-^* OT-1 T cells. Like CAR-T cells, *Prf1^-/-^* OT-1 T cells kill antigen-negative TBK1/IKKε KO targets more effectively than WT targets (Fig. S4G). Paracrine killing by OT-1 T cells was similarly dependent on TNF or IFNγ (Fig. S4G). Using OT-1 T cells co-cultured with cOVA-expressing macrophages differentiated from Hoxb8-immortalized progenitors (*23*), we demonstrated that T cells activated by an APC can also carry out paracrine killing of targets with altered TNFR1 signaling (Fig. S4H).

We next tested whether genetically mimicking the effect of a microbial-encoded TBK1/IKKε antagonist in target cells sensitized them to elimination *in vivo*. We subcutaneously implanted 2.5 × 10^5^ WT or TBK1/IKKε DKO B16 cells and 5 days later infused with PBS or *Prf1^+/+^*CAR-T cells against B7H3 (5 × 10^6^). CAR-T cell infusion did not control the growth of WT tumors whereas it delayed the growth of the TBK1/IKKε DKO tumors (Fig. S5A). Since these targets expressed B7H3, we could not conclude from this experiment that target elimination was due to contact-independent paracrine killing. To do so, we implanted into 4 groups of mice B16 tumors with the following composition: (i) 100% B7H3 KO; (ii) 90% B7H3 KO spiked with 10% WT; (iii) 100% B7H3/TBK1/IKKε TKO; (iv) 90% B7H3/TBK1/IKKε TKO spiked with 10% WT. Infusion of *Prf1^-/-^*CAR-T cells, which are defective in perforin-dependent killing, suppressed tumor growth only in cohort (iv) with B7H3/TBK1/IKKε TKO spiked with B7H3-expressing WT cells (Fig. 3E). Subsequent blotting of endpoint tumors harvested from 3 mice in cohort (iv) revealed that those tumors expressed TBK1 (Fig. S5B) and B7H3 (Fig. S5C), inferring that the TBK1/IKKε-deficient targets were selectively eliminated by the CAR-T cells whereas TBK1/IKKε-sufficient WT cells from the 10% spike survived. These genetic ablation studies strongly suggest that the presence of a pathogen-derived antagonist in target cells could result in their selective elimination during paracrine killing. Since target discrimination is imposed by the presence of a pathogen-derived factor, we propose the term ‘pathogen-restriction’ to describe this discriminatory mechanism.

### Microbial-encoded molecules function as pathogen-restriction elements

To test the pathogen-restriction hypothesis further, we examined two pathogen-encoded antagonists. YopJ, a bacterial effector from *Yersinia* pestis, acetylates the activation loop of TAK1 to block its activity and downstream NFκB and MAPK activation (*15, 24*). VP35 encoded by EBOV functions as a pseudo-substrate to inhibit TBK1/IKKε phosphorylation of IRF3/7 and type I IFN expression (*16*). ORFs encoding YopJ, VP35, or as a control hCD19, were cloned upstream of a T2A-ZSGreen cassette in an expression vector. We transiently transfected these constructs into B16 cells, which were then treated with TNF and IFNγ (10 ng/mL) for 24 hours, and then assessed viability by flow cytometry. Cytokine treatment of the VP35- and YopJ-expressing cells resulted in a reduction in their ZSG fluorescence intensity but had minimal effect on the hCD19-expressing control cells (Fig. 4A). It was also evident that there was reduced green fluorescence in the live cell gate accompanied by its increase in the debris gate on FSC/SSC plots from the cytokine-treated, YopJ- and VP35-expressing samples, indicative of extensive cell death. Blotting of lysates generated in the same manner showed activation of CASP8 and CASP3 in B16 cells expressing YopJ and VP35 treated with cytokines (Fig. S6A). Thus, presence of microbial-derived molecules confers selectivity for killing by cytokines.

**Fig. 4.**
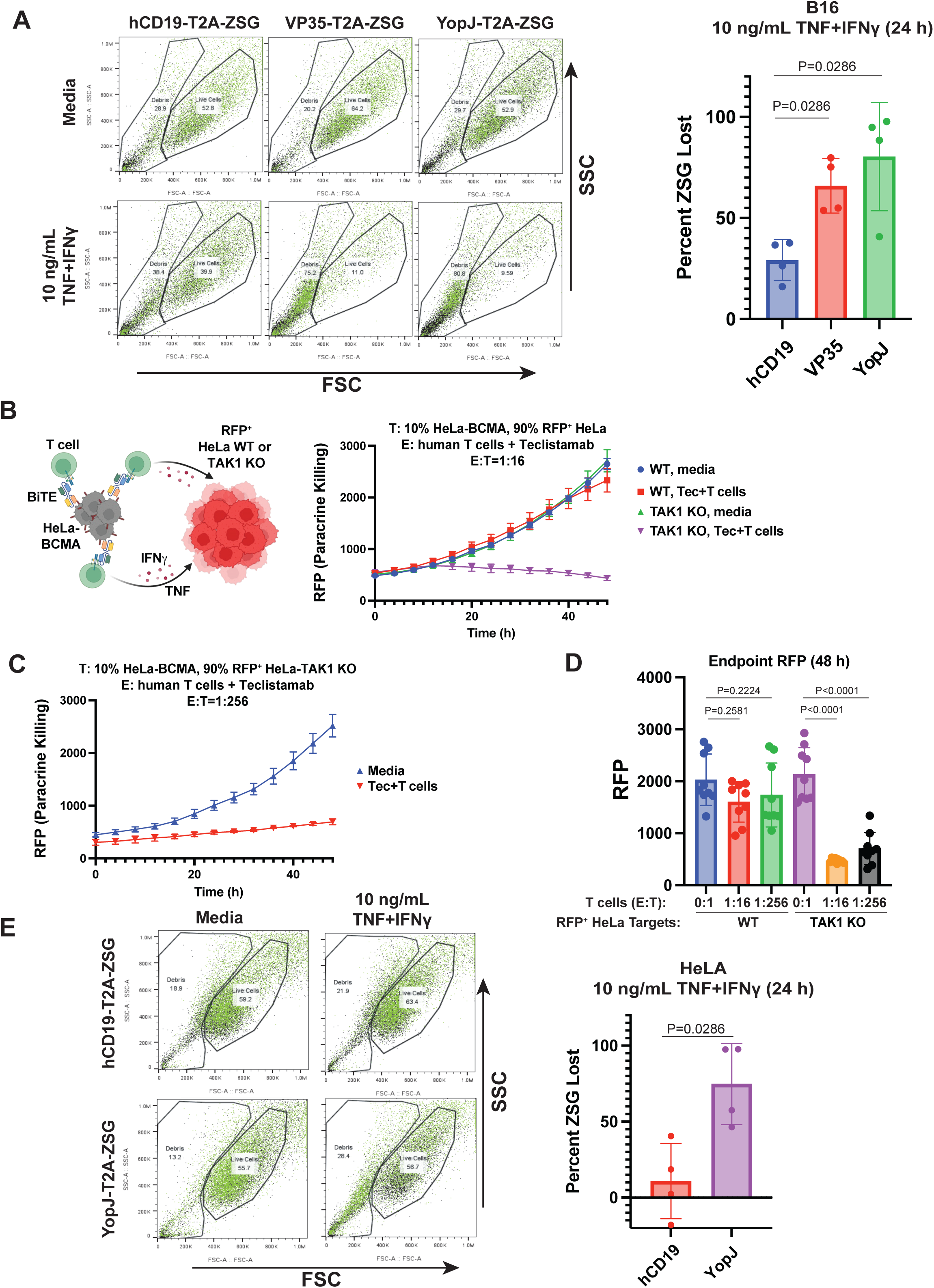
Microbial-derived factors function as restriction element during paracrine killing. **(A)** B16 cells were transiently transfected with expression plasmids encoding FLAG-VP35-T2A-ZSG, FLAG-YOPJ-T2A-ZSG or hCD19-T2A-ZSG as negative control. The next day, cells were stimulated with 10 ng/mL TNF and IFNγ for 24 hours followed by flow cytometry analysis. Dot plots are FSC versus SSC for each sample with an overlay of ZSG fluorescence in green. Representative dot plot from one experiment is shown. Bar chart shows the percentage loss in ZSG signal in cytokine-treated samples relative to media for each transfected DNA from four experiments. Values are mean ± S.D. **(B)** RFP-labeled WT or TAK1 KO HeLa cells were mixed with 10% unlabeled BCMA-transfected HeLa cells and co-cultured with healthy donor human T cells at varying E:T ratios in the presence of the BiTE teclistamab (2 ug/mL). Paracrine killing was quantified by RFP counts of the BCMA-negative HeLa targets in the IncuCyte. Data from E:T ratio of 1:16 is shown. **(C)** Data from paracrine killing of TAK1 KO HeLa targets at an E:T ratio of 1:256 is shown. **(D)** Experiment in (B) was independently performed three times. Bar graphs represent mean RFP counts in the wells at endpoint from three biological replicates. Data from E:T ratios of 1:16 and 1:256 are shown. **(E)** HeLa cells were transiently transfected with expression plasmids encoding FLAG-YOPJ-T2A-ZSG or hCD19-T2A-ZSG as negative control. The next day, cells were stimulated with 10 ng/mL TNF and IFNγ for 24 hours followed by flow cytometry analysis as described in (A). Dot plot shown is representative of four experiments that were performed. Bar chart shows the percentage loss in ZSG signal in cytokine-treated samples relative to media for each transfected DNA from four experiments. Values are mean ± S.D. *p* values from Mann-Whitney tests are shown for data in (A, D and E).

Finally, we examined whether paracrine killing is also a feature of human T cells using HeLa cells as targets. We first generated TAK1 KO HeLa cells to mimic YopJ antagonism. WT HeLa cells were completely resistant to killing by TNF and IFNγ but the TAK1 KO were highly sensitive (Fig. S6B). We then labeled WT and TAK1 KO HeLa targets with RFP and spiked them with 10% unlabeled HeLa cells transduced with BCMA. The spiked cultures were incubated with human T cells and teclistamab, a bispecific T-cell engager (BiTE) targeting BCMA used in immunotherapy of multiple myeloma (*25, 26*), and imaged in the IncuCyte. The BCMA-negative RFP-labeled TAK1 KO, but not WT targets, were efficiently eliminated by T cells and teclistamab (Fig. 4B). As seen with B16 targets, TAK1 ablation greatly sensitized HeLa targets, showing paracrine killing at a very low E:T ratio of 1:256 (Fig. 4C & 4D). Conditioned media from cultures containing BCMA-expressing HeLa cells, teclistamab and T cells also preferentially kill TAK1 KO HeLa targets (Fig. S6C). We then tested whether presence of YopJ also conferred selectivity to killing of HeLa targets by cytokines using the YopJ-T2A-ZSG construct. HeLa targets expressing YopJ became markedly sensitive to killing by combined TNF/IFNγ, as evidenced by reduction in ZSG fluorescence and increased accumulation of ZSG-labeled debris on FSC/SSC plots (Fig. 4E). We conclude that human target cells harboring a pathogen-derived factor is also selectively eliminated by paracrine killing.

## DISCUSSION

Our study establishes that T cells have two modes of killing targets that are linked but each with distinct roles. The first is the classical perforin-mediated, contact-dependent killing where target specificity is imposed by TCR recognition of the pathogen-derived peptide. The second is cytokine-mediated paracrine killing of targets where target selectivity is imposed by a pathogen-derived factor. Because of the paracrine nature of cytokines, the latter mechanism leverages the activation process of the former to enable the elimination of many more targets, in a manner that continues to be guided by the pathogen (Fig. S7). Microbial-encoded antagonists of the TNF pathway as we showed here is one mechanism of pathogen-restriction during paracrine killing. We suspect there are many different mechanisms of pathogen-restriction, some of which may be unique to a pathogen, while others may be generic and shared amongst a class of pathogens. At its core, pathogen-restriction is the outcome of a signaling crosstalk between a pathogen-derived factor with the signaling pathways of the TNF and IFN receptors, which converts the response to the cytokines from non-lethal to lethal. This crosstalk can occur at the post-translation level as we show here, or potentially at the transcription level due to the pathogen-derived factor altering the transcriptional profile of the target cell. In either case, the pathogen-derived factor is ‘sensed’ by the TNF and IFNγ signaling pathways.

Elements of our pathogen-restriction model have been previously described in different contexts. Ablation of the genes we analyzed are known to sensitize cells to TNF-induced death (*18–22*), and they and other components of the TNFR1 signaling pathway have been identified in genetic screens to play a role in tumor killing by T cells and immunotherapy (*27–29*). The ability of combined TNF and IFNγ to kill some cell lines but not others is also known (*10*), as is the capability of T cells to eliminate antigen-loss variant tumors (*30, 31*), a process referred to as bystander killing. The present study has connected these existing dots in controlled models to reveal another mechanism of T cell cytotoxicity that eliminates targets with high precision. The term bystander killing implies that the target is eliminated for no apparent reason and it is highly unlikely that our immune system would have evolved such a wasteful mechanism. Rather, we argue that cytokine-mediated killing is a highly targeted mechanism that minimizes harm to uninfected bystander cells, and our pathogen-restriction hypothesis would suggest that targets thought to be killed in a bystander manner in prior studies occurred because they harbored pathogen-restriction elements that we do not understand. We suggest that paracrine killing is a more accurate term.

The fundamental understanding that paracrine killing functions to kill target cells in a discriminatory manner provides additional insights into other aspects of our immune system. Polyfunctional T cells that co-express TNF and IFNγ have been described to be highly potent in anti-pathogen responses (*32, 33*). Our data suggest that this may be because these dual producers are capable of paracrine killing of infected and tumor targets. Because target elimination by paracrine killing does not require an MHC/Ag complex to be displayed, T cells can use this mechanism to eliminate targets with missing self-MHC, such as cells infected by viruses that downregulate MHC. This function has been thought to be mainly carried out by NK cells (*34, 35*). Since CD8, CD4 and NK cells can all produce TNF and IFNγ, they can kill targets with missing self in a paracrine manner if the targets also harbor a pathogen-restriction element.

Understanding pathogen-restriction and how paracrine killing works has significant clinical implications. Since we utilize CAR-T cells and tumor targets to understand this fundamental process, the translatability into tumor immunotherapy is clear. Mimicking pathogen-restriction in tumor targets may sensitize their response to immune checkpoint blockade, CAR-T and BiTE therapy. Our data also suggests that activating an existing pool of T cells that are not tumor-specific can have an impact on tumor targets, but this must be coupled with a strategy to mimic an infection in the tumor targets. Finally, the destructive power of paracrine killing suggest that it is also likely to play a role in T_H_1 pathologies such as irAEs, autoinflammation, GVHD and allograft rejection. A thorough understanding of pathogen-restriction mechanisms and whether these are dysregulated may provide insights into the etiology of these disorders.

## LIMITATIONS

How IFNγ synergizes with TNF to kill target cells remained unclear at the molecular level. Since IFNγ induces many genes, it is likely that multiple IFNγ-regulated genes function in concert to enhance the killing capacity of TNFR1. One candidate is TNFR1 itself as it is induced by IFNγ (GSE278131) but ablation studies with TNFR1 will not be meaningful as TNF can no longer signal. We also did not perform infection studies with Yersinia pestis and EBOV, the two pathogens encoding the TNFR1 antagonists we used as pathogen-restriction elements in our studies. The role of paracrine killing during infections and how pathogens circumvent this anti-microbial response will be of great interest. In the tug-of-war between host and pathogens, we fully expect pathogens to possess mechanisms to disarm paracrine killing. Future studies would require the use of mutant strains defective in their ability to do so.

## MATERIALS & METHODS

### Retroviral and lentiviral transduction of B16 F1 cells

To generate lentivirus, an 80% confluent plate of 293 EBNA cells were transfected with 2.5 μg Peak8-VSVG, 7.5 μg psPAX2 encoding gag-pol (Addgene, 12260), and 10 μg of lentiviral plasmid packaged with 30 μL of Lipofectamine-2000 in serum-free, plain DMEM. The following day, media was aspirated and replaced with complete media for B16 cells (DMEM, 10% FBS, 100 IU/ml penicillin and 100 µg/mL streptomycin). After two days, the virus-containing media was collected and spun in a Beckman Coulter Ultracentrifuge at 49,600 x *g* for 90 min at 4 °C. The viral supernatant was concentrated down to 1 mL and used to resuspend 1 x 10^6^ B16 cells with 4 μg/mL polybrene. The resuspended B16 cells were then plated in a six well plate, wrapped in saran wrap, and spun at 859 x *g* for 90 minutes. The cells were then placed in a 37°C incubator for two days before the media was replaced with 2 mL of B16 media together with antibiotics for selection. For generating retroviruses, the same procedure was used except that pMD.OGP (*36*) was used to express the retroviral gag-pol.

### Expression constructs

The RetroHygro vector used for stable expression was generated in-house from the Moloney murine leukemia virus (MMLV) vector pMMP412 (*37*) into which an internal ribosome entry site (IRES)-hygromycin resistance cassette was inserted downstream of the ORF of interest. ORF encoding human CD19 was amplified from an existing plasmid and cloned into an in-house derivative of pLVX-EF1alpha-SARS-CoV-2-nsp13-2xStrep-IRES-Puro (gift from Dr. Nevan Krogan, Addgene #141379) modified to encode T2A-hygromycin cassette downstream of the hCD19 insert. The hCD19 insert was replaced by a gene fragment encoding cytoplasmic ovalbumin (cOVA) to generate a cOVA expressing construct. Gene fragments encoding FLAG-tagged EBOV VP35 and FLAG-tagged YopJ were also cloned into the pLVX-EF1alpha vector upstream of a T2A-ZSGreen cassette. Retroviral constructs encoding 3^rd^ generation murine CARs were generated in the pMSGV plasmid upstream of an IRES-Thy1.1 cassette. ScFv fragments from FMC63 (against human CD19) or m276 (against murine B7H3) were fused to mCD8TM-CD28-41BB-CD3zeta. All synthetic gene fragments used for plasmid constructions were ordered from TwistBio. All inserts were verified by sequencing prior to use.

### Transient transfections

B16 or HeLa cells were plated in a 24-well tissue-culture plate at 2 x 10^5^ cells/well and incubated overnight. The following day, media was aspirated and replaced with plain DMEM containing 10 ul/mL lipofectamine-2000 and 1 ug/mL of plasmid DNA expressing hCD19, VP35, or YopJ-ZSG fused to T2A-ZSG-NLS. 24 h later, the culture media was replaced with fresh complete DMEM media (DMEM, 10% FBS, 100 IU/ml penicillin and 100 µg/mL streptomycin) and stimulated with cytokines for a further 24 h. ZSG expression was analyzed by flow cytometry.

### Generation of knockout cell lines by ribonucleoprotein (RNP) transfection

For each gene, two different target crRNAs (100 μM) were individually combined with tracrRNA (100 μM) at a 1:1 ratio, incubated at 95°C for 5 minutes and cool to room temperature to form duplexes. crRNAs and tracrRNA were synthesized by IDT. A mixture of 4.5 μL of each duplex and 4.5 μL of Cas9 enzyme (MacroLab Facility, UC Berkeley) was incubated at room temperature for 10 minutes to form RNP complex. 1 x 10^6^ B16 cells were resuspended in Lonza SF nucleofection buffer and combined with the RNP complex. Cells were electroporated using Lonza 4-D nucleofector pulse code DJ-110. HeLa KOs were prepared the same with the pulse code (CN-114). Western blotting or flow cytometry was carried out 48-72 hours post electroporation to analyze the efficiency of gene ablation. To generate stable knockouts for long term studies, the nucleoporated cells were single cell cloned by limiting dilution and multiple clonal lines screened for gene knockout by western blotting or flow cytometry. Multiple clones with verified knockout were then pooled for use to minimize potential founder effects.

### Target sequences of sgRNA for RNP-mediated knockout

CRISPR targets for gene knockouts were identified using CRISPick (Broad Institute).

*Tbk1* (TBK1): ACGGGGCTACCGTTGATCTG and TTTGAACATCCACTGGGCGA

*Ikbke* (IKKe): GGACAATGCCATTCTCCCGC and CCATGATTAGCACCTTCTGC

*Rnf31 (*HOIP): CAGCCGAGGTCGCTCACATA and ACAGCTCTGAACGCAACCAC

*Map3k7* (TAK1): GTGCAGGTAAGCCACTCCTT and AAACCCTTCGATGAGATCGG

*Cd276* (B7H3): GGTCATGCTGGGCTTCGAGT and GAGCGCTGTGCGGTTGGAGT

*Fadd* (FADD): GGCGCTGCAGTAGATCGTGT and CCTGTCGGGCAACGATCTGA

*Stat1*(STAT1): GGTCGCAAACGAGACATCAT and GGTCGCAAACGAGACATCAT

*MAP3K7* (TAK1): GCGTGGGCAGCAGTATAATA and CATGTGTGTCTGAATGTCAC

### Generation of CAR retroviruses

Ecotropic retroviruses encoding murine CARs were generated in the same manner as described above except using 5 µg pCL-EcO to code for gag-pol and envelope. 24 h post-transfection of 293EBNA packaging cells, media was replaced with complete splenocyte media (RPMI 1640, 10% FBS, 1X Pen-Strep, 1% L-glutamine, 1% non-essential amino acids, 1% sodium pyruvate, and 0.1% beta-mercaptoethanol). After another 24 h, the viral supernatant was harvested, centrifuged at 483 x g for 5 minutes to pellet cellular debris and used to resuspend splenocytes.

### T cell generation from splenocytes

Spleens were harvested from C57BL/6 WT and *Prf1-/-* mice and manually homogenized using a syringe plunger. The homogenate was filtered through a 40 µm strainer with sterile PBS and pelleted by centrifuging at 1200 RPM for 5 minutes. The pellet was treated with 3 mL of ACK buffer (Gibco A1049201) for 90 seconds to lyse red blood cells. ACK-treated splenocytes were suspended in 12 mL of complete splenocyte media and pelleted. Cells were resuspended in 10 mL of complete splenocyte media and counted on an Invitrogen Countess II. Cells were then plated in a 24-well tissue-culture plate (Falcon 353047) at 2 × 10^6^ cells/mL and treated with Concanavalin A (ConA, 0.25 µg/mL) and murine IL-2 (1 µg/mL) overnight.

### Retroviral transduction of T cells

One day prior to the transduction, a 24-well non-tissue culture plate (CELLTREAT 229524) was coated with 0.5 mL/well of RetroNectin (Takara Bio T100B, 25 µg/mL in PBS) and stored overnight at 4°C. The following day, the RetroNectin coated plate was aspirated and blocked with 2% BSA in PBS (0.5 mL/well) for 30 min. The BSA solution was aspirated and the wells washed twice with 1 mL of sterile PBS. ConA/IL-2-treated T-cells were counted and resuspended in 1 mL of viral supernatant per 10^6^ T-cells. Murine IL-2 (50 U/mL) and polybrene (0.8 µL/mL) were added to the T-cells/viral mixture. Cells were dispensed into the RetroNectin coated wells at 1 mL/well and the plated sealed with parafilm. This plate was centrifuged at 800 × g for 90 min at 32°C. After the spinoculation, 1 mL of splenocyte medium supplemented with ConA and IL-2 was added to each well and incubated overnight. The following day, cells were split 1:2 into fresh tissue-culture 24-well plates and 1 mL of splenocyte medium supplemented with ConA and IL-2 was added to each well. After two days of incubation, CAR T cells were validated by flow cytometry for Thy1.1 and were maintained with IL-2 supplementation every 3 days. CAR-T cells were optimal for functional assays on days 7–8 post-transduction.

### Human T cell cultures

Apheresis cones (*38*) were obtained from the Mayo Clinic Blood Donor Center and Ficoll separated into peripheral blood mononuclear cells (PBMCs). PBMCs were then cryopreserved in FBS + 10% DMSO. Upon thawing, PBMCs were magnetically sorted into purified T cells using the EasySep Human T Cell Isolation Kit from STEMCELL Technologies (17951). T cells were stimulated with soluble anti-CD28 (Biolegend 302934) and anti-CD3 (Biolegend 300438) at 10 ug/mL. Media was supplemented with rhIL-2 (PeproTech #200-02) starting at 200 IU/mL and coming down to 50 IU/mL in a stepwise fashion over 5 days.

### Western blotting

Whole-cell lysates were obtained using triton lysis buffer (20 mM Tris-HCl pH 7.4, 40 mM NaCl, 5 mM EDTA, 50 mM NaF, 30 mM Na Pyrophosphate, 1% Triton X-100) that contained 1x protease inhibitor (Millipore Sigma, 539137) and 1x phosphatase inhibitor (Thermo Scientific, 78426). Protein concentration was measured using Pierce BCA (Thermo Scientific, 23227). 50 or 100μg of protein samples were boiled at 95°C in 1X SDS sample buffer and resolved by reducing SDS-PAGE. Resolved proteins were transferred to nitrocellulose membranes (Amersham, 10600003), blocked with 5% milk in 1X TBST solution for 1 hr at room temperature, followed by overnight incubation with primary antibodies at 4°C. After a series of washes with 1X TBST, membranes were incubated with secondary HRP-conjugated antibodies in 2.5% milk in 1X TBST solution. After multiples washes, membranes were incubated in chemiluminescent substrate solution (Thermo Scientific, 34076) for 2 minutes and imaged with the BIO-RAD ChemiDoc MP instrument. For reblotting, membranes were stripped with guanidine HCl, blocked with milk and reprobed with subsequent antibody.

### Nuclear Extract Preparation

Nuclear extracts were prepared by plating 5 × 10^6^ in 10 cm plates and stimulated with cytokines overnight. The following day, cells were harvested by scraping, washed once with ice-cold PBS, and pelleted by centrifugation at 14,000 rpm for 30 sec. Cells were resuspended in 400 μl Buffer A (10 mM HEPES, pH 7.9; 10 mM KCl; 0.1 mM EDTA; 0.1 mM EGTA) supplemented with 1 mM DTT, 1 mM PMSF, and protease inhibitors. This was incubated on ice for 15 min. NP-40 (10%, 25 μl) was added, and samples were vortexed vigorously for 10 s, followed by centrifugation at 14,000 rpm for 1 min. The supernatant was removed, and the nuclear pellet was washed once with Buffer A containing inhibitors. Nuclei were extracted in 50 μl Buffer C (20 mM HEPES, pH 7.9; 0.4 M NaCl; 1 mM EDTA; 1 mM EGTA) supplemented as above, and samples were shaken at maximum speed for 30 min at 4°C. After centrifugation (14,000 rpm, 1 min), the supernatant containing nuclear proteins was collected and stored at −80°C. Protein concentration was measured using Pierce BCA.

### Cell death quantification

Target tumor cells were seeded at 4000 cells/well in a 96-well tissue culture plate. After overnight culture, the old media was replaced with fresh complete media containing recombinant cytokines, CAR-T cells, or OT-1 T-cells at varying concentrations, together with 0.5 μM of the cell-impermeable viability dye YOYO-3 (Thermo Scientific, Y3606). The cultures were analyzed using a Sartorius IncuCyte S3 live-cell imaging system and four images of each well were taken every 4 hours. Cell death events were quantified as a measure of YOYO-3 fluorescence counts normalized to the confluency at each time point. For paracrine killing analysis, target cells were labeled with the raspberry fluorescent protein (RFP) and RFP signals quantified by the IncuCyte software. For paracrine killing by human T cells, we mixed unlabeled HeLa cells transfected with BCMA (HeLa-BCMA) with RFP-labeled HeLa targets, which do not express BCMA. T cells from healthy human donors were added together with 2 ug/mL of teclistamab, a BiTe specific for BCMA and CD3. Paracrine killing was assessed by loss of RFP signal in an IncuCyte S3 instrument. Cell death analyses in the IncuCyte are typically conducted with varying titrations of cytokines or with T cells in a range of E:T ratios. For simplicity, data from a single dose or E:T ratio is generally shown.

### APC:T cell:target co-cultures

ER-HoxB8 Cas9-GFP murine marrow progenitor (MMP) cells [provided by Dr. David Sykes (Massachusetts General Hospital)] were cultured in DMEM supplemented with 10% FBS, 100 IU/ml penicillin, 100 µg/mL streptomycin, 10 ng/mL GM-CSF, and 0.5 uM beta-estradiol. The MMP cells were transduced with the lentivirus encoding cOVA described earlier and selected with 500 ug/mL hygromycin. To differentiate MMPs to macrophages, 1x 10^6^ cells were washed twice in PBS and seeded in a 96-well plate for 24 hours at 400 cells/well in same media without beta-estradiol. 9-fold more RFP-labeled B16 targets were then added to the wells, followed by the addition of OT-1 T cells at varying E:T ratios. Paracrine killing was assessed by loss of RFP signal in an IncuCyte S3 instrument.

### *In vivo* tumor studies

All animal studies were carried out in accordance with approved IACUC protocol at the Mayo Clinic in Rochester, MN. C57BL/6 (Jax #664), C57BL/6-Tg (TcraTcrb) 1100Mjb/J (OT-I) (Jax #3831) and *Prf1-/-* (JAX #002407) were obtained from Jackson Laboratory and housed in standard mice rooms. Tumor challenge experiments were performed with mice 8 weeks or older. Mice were shaved at the inoculation site a day before tumor implantation. 2.5 x 10^5^/100 μL B16 F1 tumor cells were resuspended in sterile PBS (Corning) and subcutaneously injected into the right flank on day 0. Prior to all tumors becoming palpable (day 5), mice were randomized for the single treatment of either PBS (100 μL) or generated CAR-T cells (5 x 10^6^/100 μL) through intravenous injection via tail vein. Tumor volume was measured every 2-3 days using a digital caliper until either survival end point was reached or till the end of study. Tumor end points were adhered to as defined by the IACUC protocol. Mice were euthanized by AVMA-approved CO_2_ asphyxiation. At least five mice were used in each group for all experiments.

### Enzyme-Linked Immunosorbent Assay (ELISA)

B16 melanoma cells were seeded in a 6-well plate at 2.5 x 10^5^ cells per well and stimulated for 24 h. Culture supernatants were then harvested and stored at -80°C. Samples were diluted to fall within the range of the standard curve. ELISA kit for mouse CCL2 (MCP-1) was obtained from BioLegend (BioLegend 432704) and mouse CXCL9 (MIG) was obtained from Bio-Techne (R&D DY492-05). ELISAs were performed according to manufacturer’s instructions and read on a MultiSkan SkyHigh (Thermo Fisher) plate reader. Data analysis was performed using GraphPad.

### Flow Cytometry

Cells were harvested from plates using trypsin-EDTA (Corning 25-053) and resuspend in PBS and washed three times in PBS. Subsequently, pellets were resuspended in 100 ul of FACS buffer (PBS containing 5% FBS, 0.5% BSA, and 0.1% sodium azide) with 1 ul of conjugated antibodies and stained for 30 minutes in the dark at 4°C. If primary antibody was unconjugated, the cells were washed once, and resuspended with 100 ul FACS buffer and 1 ul of secondary antibody for an additional 30 minutes in the dark at 4°C. Following surface staining, samples were washed twice with FACS buffer and data acquired using the CyTek Northern Lights Spectral Flow Cytometer (Cytek Biosciences).

### NanoString

B16 melanoma cells were seeded in a 6-well plate at 2.5 x 10^5^ cells per well. The following day, 10 ng/mL IFNγ and/or TNF were added to the wells. Cells were treated for 24 hours prior to mRNA isolation with DNase digestion (Qiagen #74106, #79254). RNA hybridization was performed for 24 h at 65°C following manufacturer recommendations (NanoString nCounter Host Response Panel). The sample cartridge was loaded by the nCounter Prep Station and ran on the nCounter Digital Analyzer. Raw counts were normalized to library size and transformed to logCPM with limma-voom (*39, 40*). Normalized counts were then used to arrange genes into clusters (*41*) and visualized with ComplexHeatmap (*42*).

### Reagents and Antibodies

Cytokines and reagents were obtained from the following sources: murine TNF (PeproTech, 315-01A), murine IFNγ (PeproTech, 315-05), human TNF (PeproTech 300-01A), human IFNγ (Thermo Scientific 300-02), murine IL-2 (PeproTech 212-12), human IL-2 (PeproTech 200-02) and Concanavalin A (Sigma-Aldrich C5275). Antagonist antibodies used were from following sources: murine-TNF (clone XT3.11, Bio X cell BE0058) murine IFNγ (clone XMG1.2, Bio X Cell BE0055), human-TNF (Bio X Cell SIM0001) and human IFNγ (Bio X Cell BE0235). For western blotting, primary antibodies used were TBK1 (Cell Signaling, 3013S), IKKε (Cell Signaling, 2690S), phospho-STAT1 (Tyr701) (Cell Signaling, 9167S), STAT1 (Cell Signaling, 9172S), cleaved caspase 8 (Cell Signaling, 8592S), cleaved caspase 3 (Cell Signaling, 9661S), cleaved PARP (Cell Signaling, 9541S), FADD (Abcam, ab124812), NFκB p65/RelA (Cell Signaling, 8242S), NFκB p105/p50 (Cell Signaling, 12540S), NFκB p100/p52 (Cell Signaling, 52583S), RelB (Cell Signaling, 4922S). β-Actin (Cell Signaling, 3700S) was used as a loading control for whole cell lysates, while HDAC1 (Cell Signaling, 5356T) was used as a loading control for nuclear lysates. Primary antibodies were used at a 1:1000 dilution in antibody buffer containing 2.5% BSA, 0.05% sodium azide in 1x TBST. Secondary antibodies against rabbit IgG (Jackson ImmunoResearch, 111-035-144) and mouse IgG (Jackson ImmunoResearch, 115-035-146) were used at 1:5000 in 2.5% milk in 1x TBST. For flow cytometry, primary antibodies were CD90 (Thy1.1, BioLegend 202503), CD276 (B7H3 Clone EPNCIR122, Abcam AB134161), and Live/dead Violet (Thermo Scientific L34963). Secondary goat-anti-rabbit (Invitrogen A11034) was used to detect the B7H3 primary antibody. Teclistamab was generously provided by Dr. Wilson Gonsalves (Mayo Clinic).

### Statistics

We performed statistical analysis using Prism GraphPad software version 10. For cell death and RFP data generated using the IncuCyte, a two-tailed Mann-Whitney U test was used to compare the endpoint data between the two groups across three biological replicates. For mouse tumor volume data, we used a two-tailed Mann-Whitney U test to compare the endpoint tumor volume between mice of different groups across all biological replicates. *p* < 0.05 was considered significant.

## Acknowledgements

We thank Dr. Christian Pfaller (Mayo Clinic) for providing HeLa cells, Dr. Miriam Merad (Icahn School of Medicine at Mount Sinai) for providing B16 F1 cells, and Drs. Yogesh Chawla and Wilson Gonsalves (Mayo Clinic) for providing teclistamab. We thank Dr. Eddie Ip (Mayo Clinic) and Dr. David Sykes (MGH) for providing the ER-HoxB8 Cas9-GFP cells generated by Dr. Sykes. We thank Dr. Sylvie Girard (Mayo Clinic) for use of the MultiSkan plate reader. We would also like to thank Dr. Brad St. Croix (NIH/NCI) for generously sharing the sequence for the m276 scFV against B7H3.

## Funding

Mayo Clinic (ATT, DDB, KAK, LRP)

National Institutes of Health (NIH) grant CA270380 (ATT)

National Institutes of Health (NIH) T32 Training Program in Immunology AI170478 (MCG, NDS, MZ), and Multidisciplinary Predoctoral Training in Discovery and Translational Research DK124190 (JM).

## Author contributions

Conceptualization: MCG, ATT

Methodology: MCG, ARC, ENK, NDS, MZ, RG, MJM, NRT, LRP

Investigation: MCG, SAR, ARC, ENK, MZ, JM, NRT

Visualization: MCG, ARC, ENK, JM

Funding acquisition: ATT, KAK

Project administration: ATT

Supervision: ATT, KAK, DBB

Writing – original draft: MCG, ATT

Writing – review & editing: MCG, AC, JM, LRP, DDB, KAK, ATT

