## Supplementary Data for "T cells use TNF and IFNγ for paracrine killing with target discrimination programmed by pathogen-derived factors"

### Supplementary Figure 1

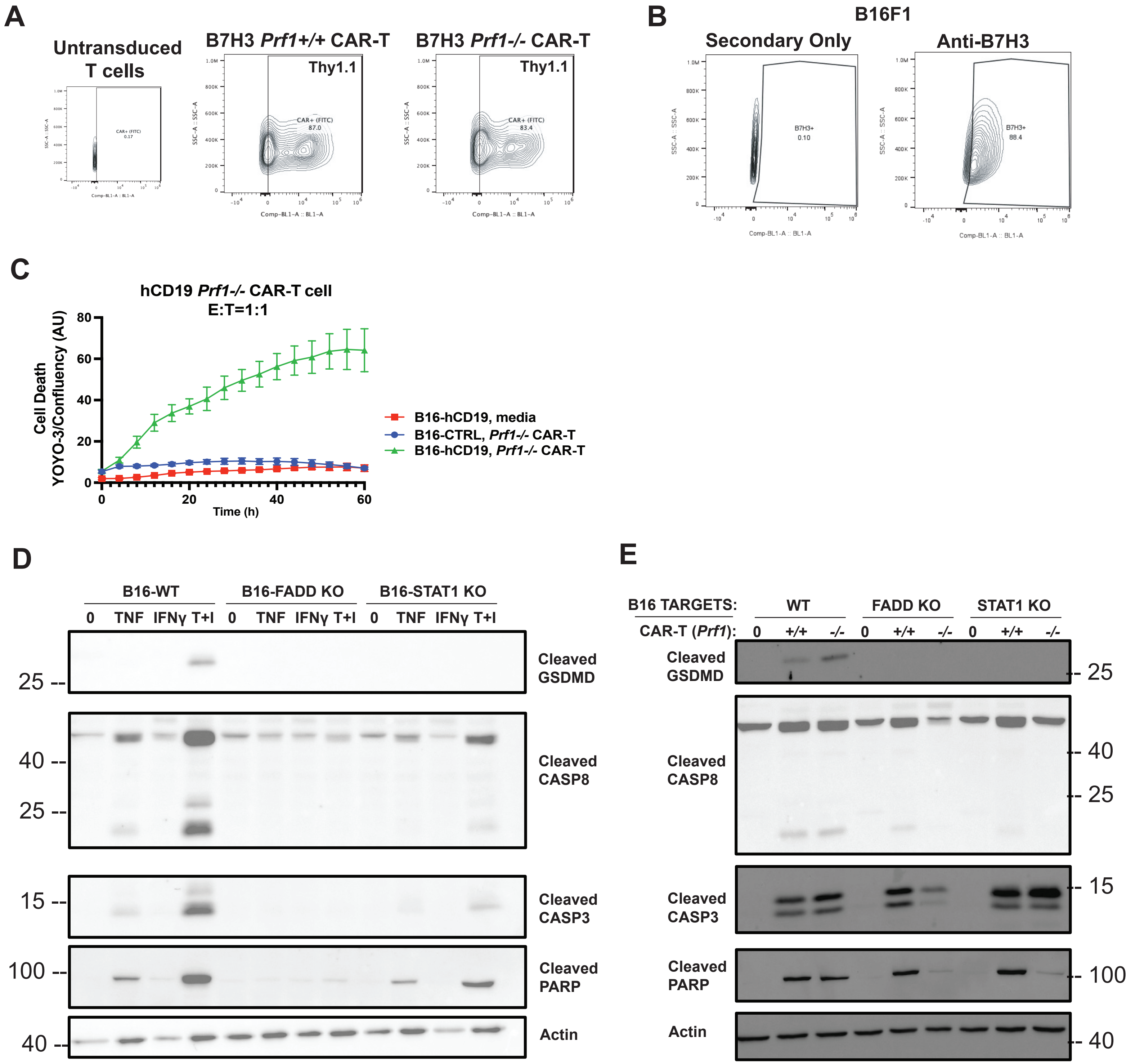

**Fig. S1. CAR-T cells validation and biochemical analysis of cytokine-mediated killing**

**(A**) Flow cytometry of *Prf1+/+* or *Prf1-/-* CAR-T cells against B7H3 (CD276) stained with anti-Thy1.1. The retroviral CAR construct consists of the m276 scFv fused to mCD8TM-CD28-41BB-CD3zeta upstream of an IRES-Thy1.1 cassette. **(B)** Flow cytometry analysis of B7H3 expression on B16 F1 melanoma cells. **(C)** *Prf1-/-* CAR-T cells against human CD19 (E:T of 1:1) were co-cultured with B16 cells stably expressing hCD19 or control protein. **(D)** Immunoblots for indicated proteins from B16 WT, FADD KO and STAT1 KO cells treated for 24 h with 100 ng/mL TNF, IFN$\gamma$, or both. **(E)** Immunoblots for indicated cell death markers from B16 WT, FADD KO and STAT1 KO cells co-cultured for 24 h with media, *Prf1+/+* and *Prf1-/-* CAR-T cells at E:T ratio of 1:1.

**Supplementary Figure 2**

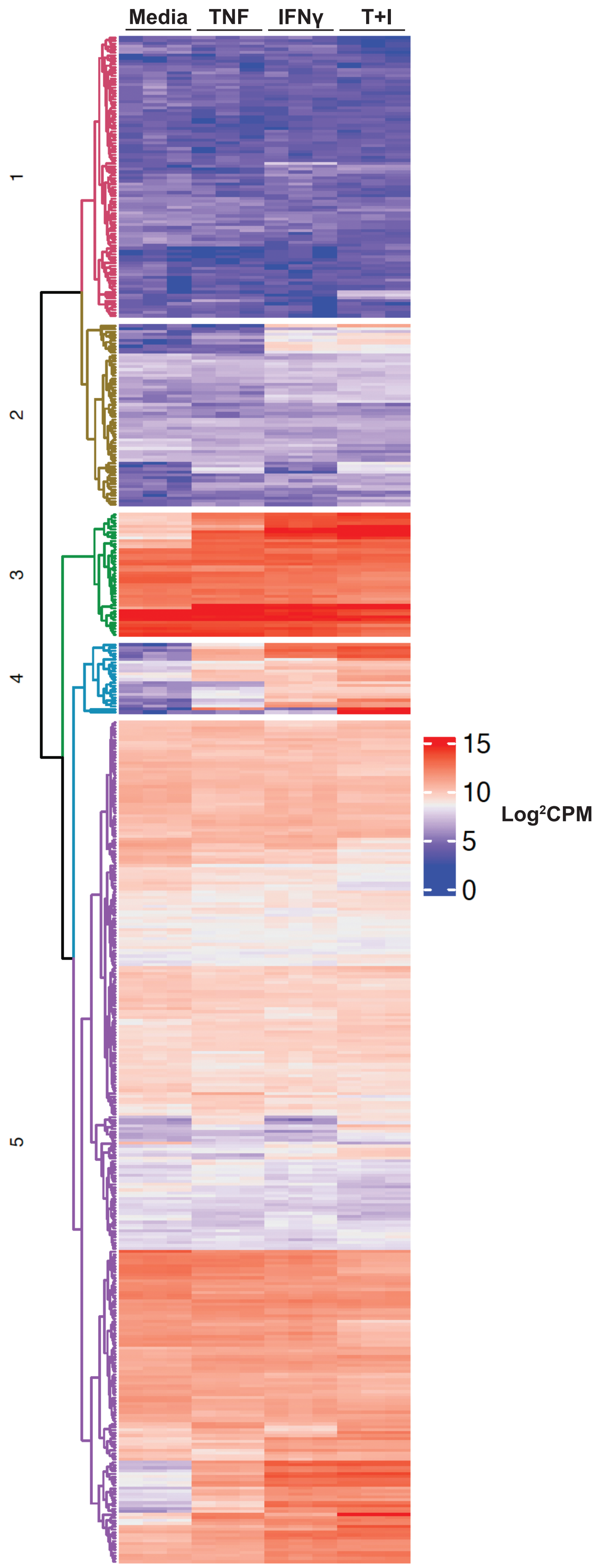

**Fig. S2. Heatmap of gene expression of B16 cell stimulated with cytokines.**
B16 WT cells were stimulated with 10 ng/mL TNF, IFN$\gamma$, or both for 24 h. RNA was isolated from three independent experiments and analyzed on the Nanostrings nCounter platform using their mouse Host Response Panel containing 775 genes. Unsupervised clustering was performed. Enlarged cluster 4 is shown in Fig. 2B. Scale indicates $Log^2$CPM.

### Supplementary Figure 3

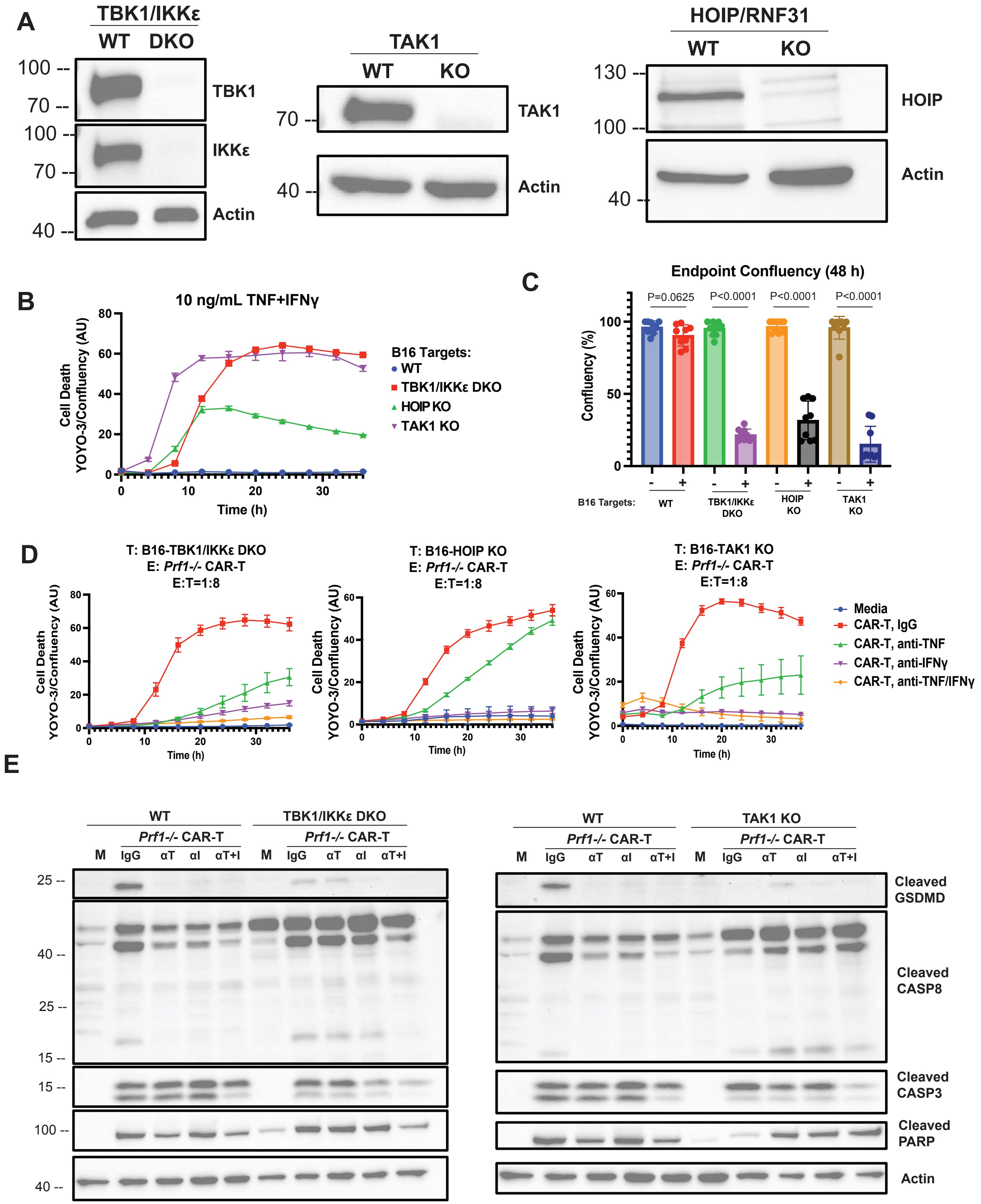

**Fig. S3. Sensitization of B16 targets to killing by cytokines and CAR-T cells.**
**(A)** RNP-mediated CRISPR knockout of TAK1, HOIP, or TBK1/IKKε in B16 cells confirmed by immunoblot. **(B)** B16 WT, TAK1 KO, TBK1/IKKε DKO and HOIP KO were treated with 10 ng/mL TNF and IFNγ and analyzed in the IncuCyte for 48 h. YOYO-3 counts normalized to confluency in each well is shown in arbitrary units. Values are triplicate mean ± SD. **(C)** The experiment in (B) was independently performed 3 times. The mean confluency at 48 h of 9 wells from the 3 biological replicates is shown in the bar chart. **(D)** B16 targets with the indicated gene knockouts were co-cultured with varying E:T ratios of *Prf1-/-* CAR-T cells. 50 ug/mL control IgG, anti-TNF, anti-IFNγ or both were added to the co-cultures and imaged. Data from E:T ratio of 1:8 is shown. YOYO-3 counts normalized to confluency in each well is shown in arbitrary units. Values are triplicate mean ± SD. **(E)** Immunoblots for indicated cell death markers of TBK1/IKKε DKO and TAK-1 KO targets co-cultured with *Prf1-/-* CAR-T cells for 24 h in the presence of IgG, anti-TNF, anti-IFNγ or both. M indicates cells treated with media. Blots shown are representative of two experiments that were performed. *p* values from Mann-Whitney tests are shown for data in (C).

### Supplementary Figure 4

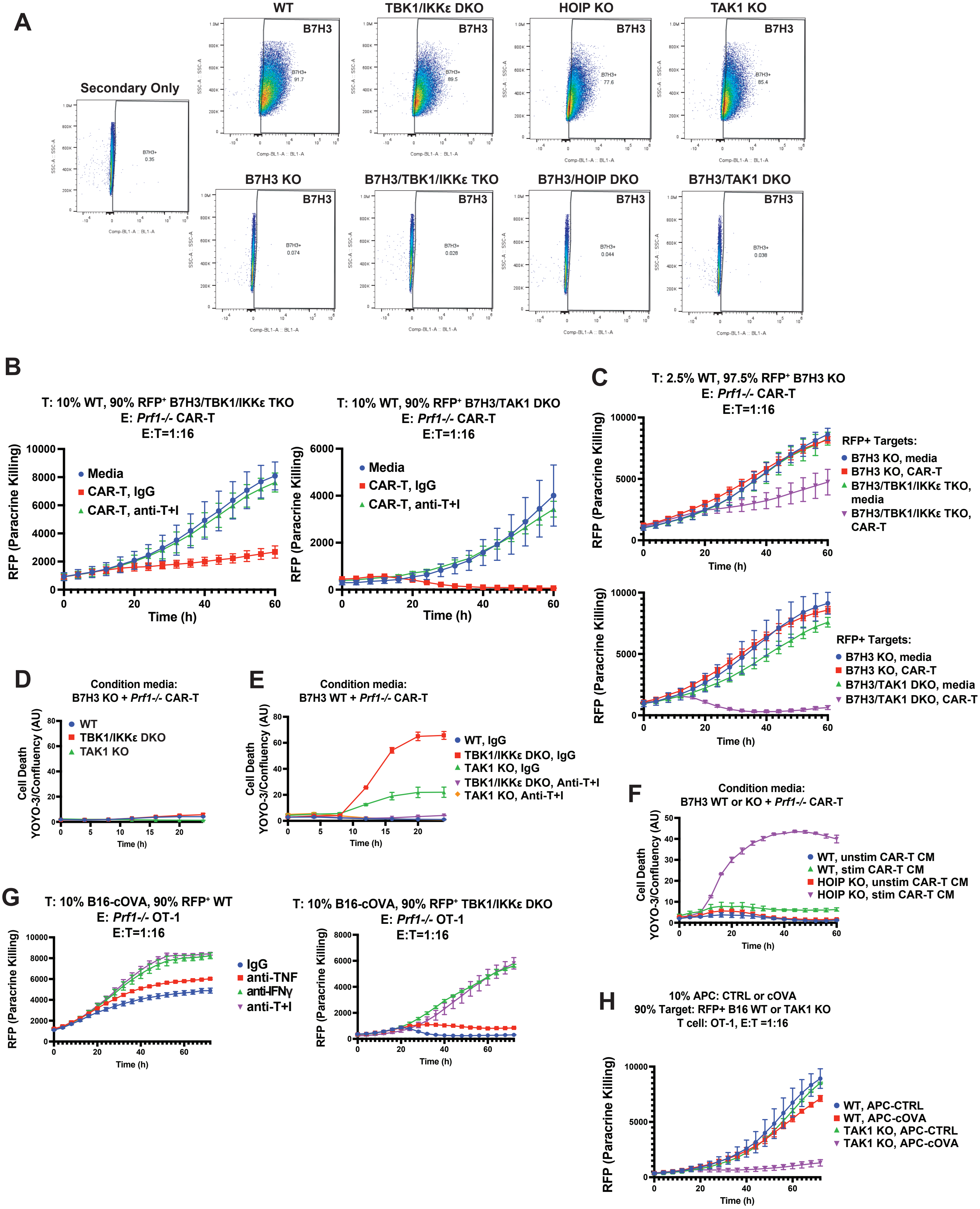

**Fig. S4. Paracrine target killing can be enhanced by mimicking the effects of microbial antagonists.**

**(A)** Flow cytometry analysis of surface B7H3 expression on B16 cells with the indicated knockouts to confirm deletion of B7H3. **(B)** RFP-labeled B7H3/TBK1/IKKε TKO or B7H3/TAK1 DKO targets were mixed with 10% unlabeled B7H3$^+$ B16 WT cells and co-cultured with *Prf1-/-* CAR-T cells at varying E:T ratios. Control IgG or anti-TNF/IFNγ (50 ug/mL) was also added to the cultures. Paracrine killing was quantified by RFP counts of the B7H3-negative targets in the IncuCyte. Data from E:T ratio of 1:16 are shown. **(C)** Lower amounts of B7H3-positive B16 cells used to spike cultures of B7H3-negative TBK1/IKKε and TAK-1 KO targets was tested. Data shown is from the lowest amount tested at 2.5%. Paracrine killing by *Prf1-/-* CAR-T cells at an E:T of 1:16 is shown. **(D)** Conditioned media from *Prf1-/-* CAR-T cells co-cultured with B7H3-deficient B16 cells were used to treat WT, TBK1/IKKε DKO and TAK1 KO targets and imaged in the IncuCyte for 24. **(E)**. Conditioned media from *Prf1-/-* CAR-T cells co-cultured with B7H3-expressing B16 cells were used to treat TBK1/IKKε DKO and TAK1 KO targets in the presence of IgG or anti TNF/IFNγ (50 ug/mL). Cultures were imaged in the IncuCyte for 24 h. **(F)** Conditioned media from *Prf1-/-* CAR-T cells co-cultured with B7H3-deficient or B7H3-sufficient B16 cells were used to treat WT or HOIP KO targets and imaged in the IncuCyte for 60 h. **(G)** RFP-labeled WT or TBK1/IKKε DKO targets were mixed with 10% unlabeled B16-cOVA cells and co-cultured with *Prf1-/-* OT-1 cells at varying E:T ratios. Control IgG, anti-TNF, anti-IFNγ, or both were also added. Paracrine killing was quantified by RFP counts of the OVA-negative targets in the IncuCyte. Data from E:T ratio of 1:16 is shown. **(H)** RFP-labeled WT or TAK1 DKO targets were mixed with 10% control or OVA-expressing macrophages and co-cultured with OT-1 cells at varying E:T ratios. Paracrine killing was quantified by RFP counts of the OVA-negative targets in the IncuCyte. Data from E:T ratio of 1:16 is shown and is representative of three experiments that were performed.

**Supplementary Figure 5**

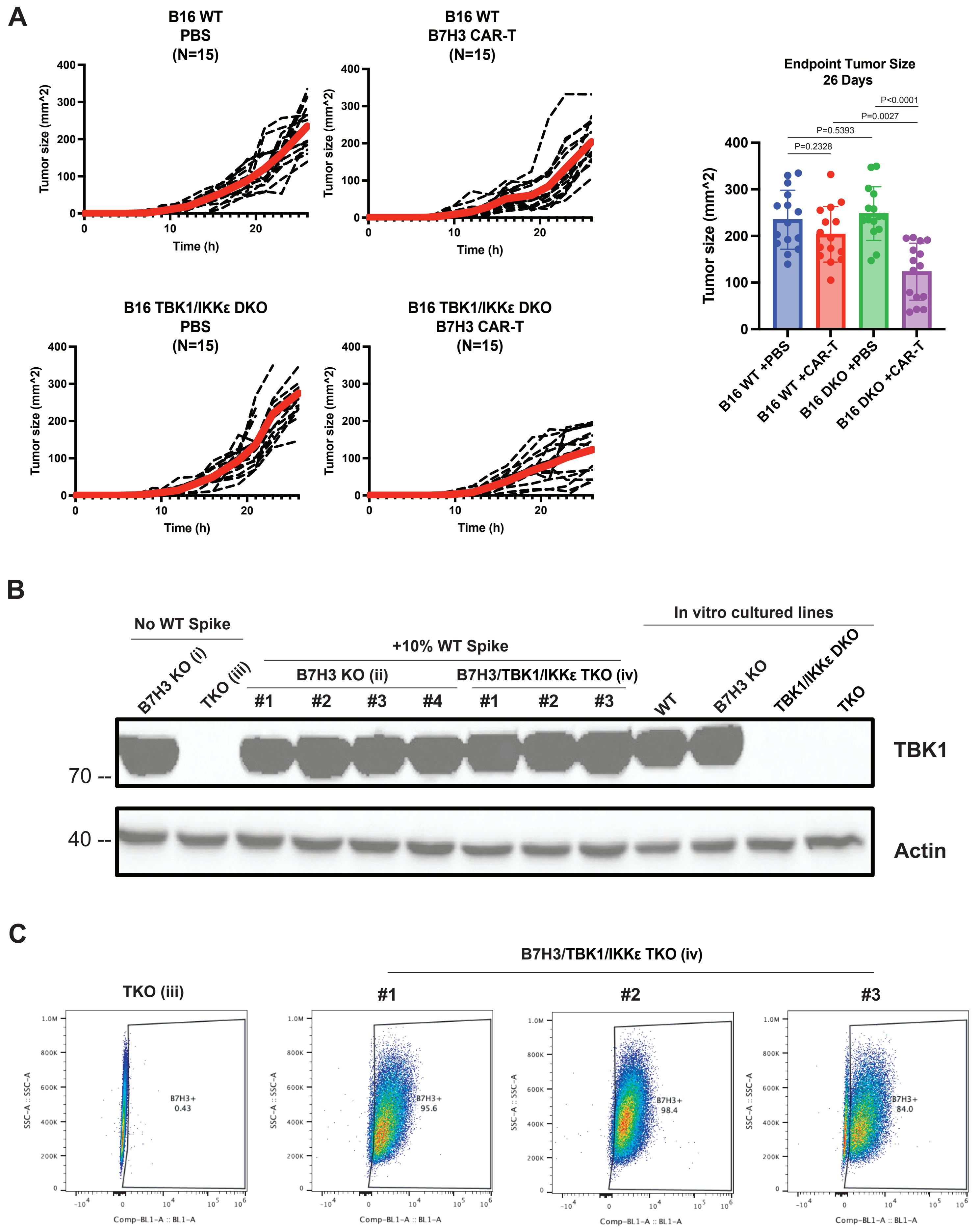

**Fig. S5. CAR-T cells control of TBK1/IKKε DKO tumors.**

**(A)** 2.5 x $10^5$ WT or TBK1/IKKε DKO B16 cells were implanted into B6 hosts. 5 days later, mice were administered PBS or *Prf1+/+* CAR-T cells against B7H3 via tail vein infusion and tumor growth measured. Spider plots shown were compiled from three independent experiments, each with N=5 per group. Thick red line indicates average tumor volume in each group of mice. Bar chart shows mean tumor volume ± SD for each group at day 26. *p* values from Mann-Whitney tests are shown. **(B)** Tumors were harvested from mice in Fig. 3E at endpoint and expanded *ex vivo*. These tumors were lysed and blotted for TBK1. The Roman numerals correspond to the groups indicated in Fig. 3E. The four lanes on the right are control lysates from the indicated cell lines grown *in vitro*. **(C)** Tumor cells from (B) were also stained with anti-B7H3 and analyzed by flow cytometry.

### Supplementary Figure 6

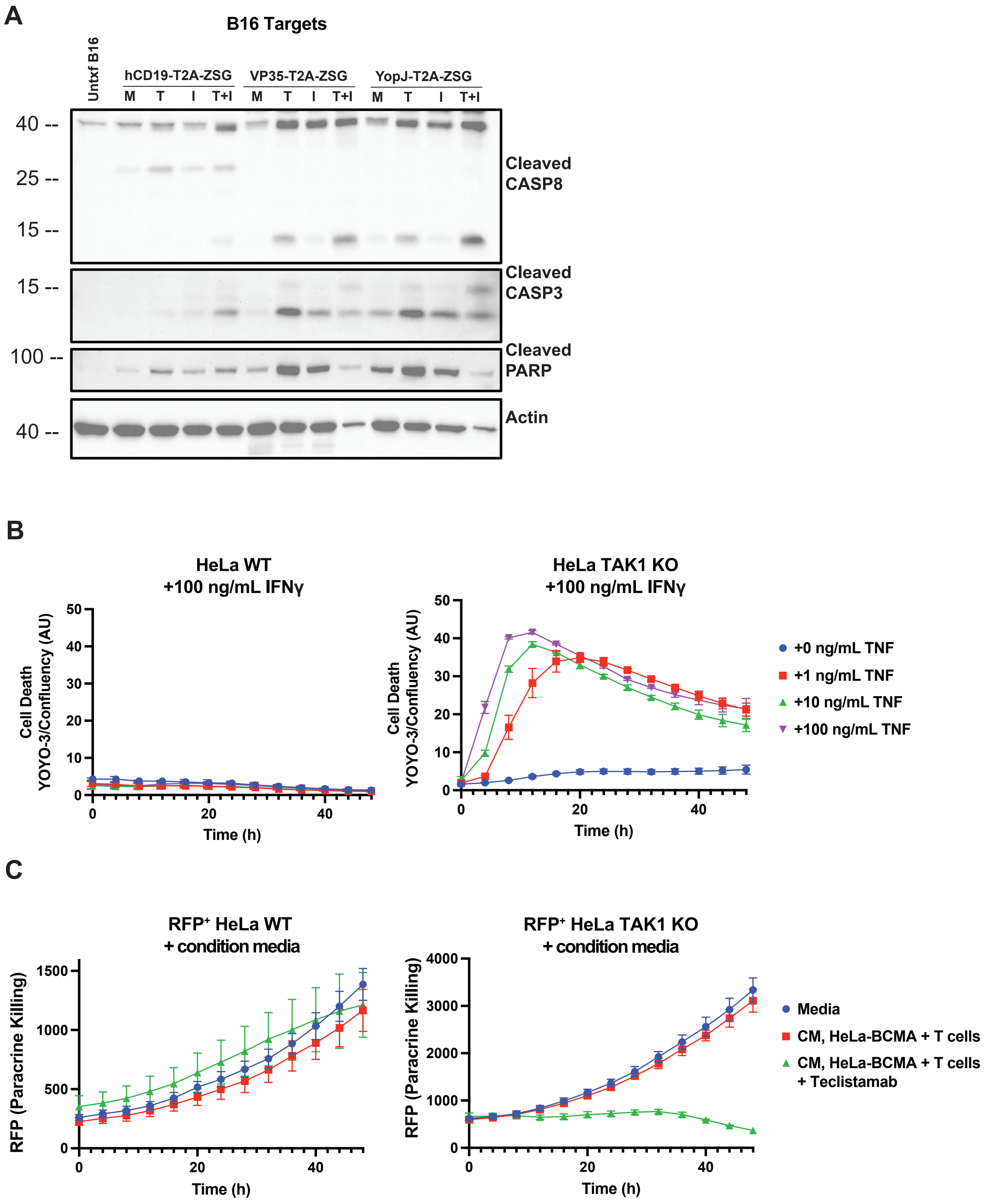

**Fig. S6. Human target cells also undergo paracrine killing by T cells.**
**(A)** Immunoblots of B16 cells transfected with hCD19-T2A-ZSG, FLAG-VP35-T2A-ZSG or FLAG-YOPJ-T2A-ZSG and stimulated with media, 100 ng/mL TNF, IFN$\gamma$ or both for 24 h. Lysates were blotted for apoptotic markers. **(B)** Hela WT or TAK1 KO cells were stimulated with increasing doses of TNF (0-100 ng/mL) in the presence of 100 ng/mL IFN$\gamma$. Target cell death was assessed by YOYO-3 staining and imaging for 48 h in the IncuCyte S3. Confluency of the cells in each well were also quantified. Data is presented as YOYO-3 counts normalized to confluency in each well in arbitrary units. Values are triplicate mean $\pm$ SD. Data from one representative experiment is shown. **(C)** RFP-labeled WT or TAK1 KO HeLa cells were treated with media or conditioned media from HeLa-BCMA cells co-cultured with healthy donor human T cells in the absence or presence of teclistamab (2 ug/mL). Paracrine killing was quantified by RFP counts of the BCMA-negative HeLa targets in the IncuCyte S3.

Supplementary Figure 7

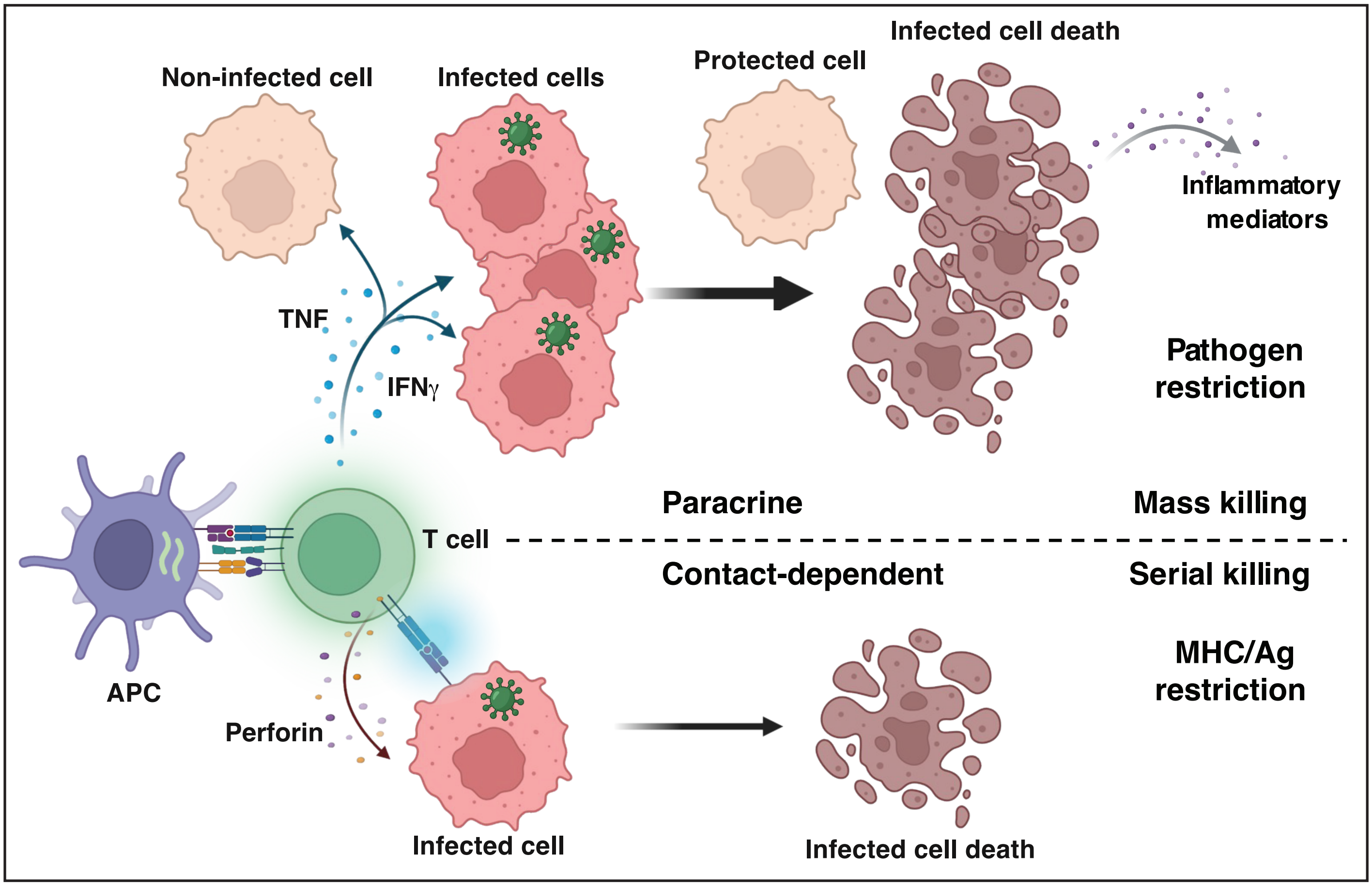

**Fig. S7. Model for pathogen-restricted paracrine killing by T cells**

T cells have two complementary modes of killing. The classical perforin-dependent killing occurs soon after TCR engagement with the MHC/Ag complex on the target cell leading to deposition of cytotoxic granules. Specificity for target killing is imposed by pathogen-derived peptides present only in the infected cell. This mechanism requires effector-target contact, occurs in a serial manner, and is likely inefficient. T cells can also carry out paracrine killing of remote targets via their secretion of diffusible TNF and IFN$\gamma$. In the default state (uninfected cell), the cytokines are non-lethal. In an infected cell, the presence of a pathogen-derived factor alters the TNF and IFN$\gamma$ signaling pathways and the outcome is now lethal. Thus, paracrine killing of targets is restricted to those harboring the pathogen, can be carried out *en masse*, and concurrently induce an inflammatory gene program in the dying cell. The T cell can be activated by an infected cell or by an APC that has taken up parts of the pathogen. The latter also allows more antigen-specific T cells to be activated, without relying solely on the infected cells. In either case, pathogen-restricted paracrine killing enables a T cell to leverage its interaction with one antigen-bearing cell to eliminate a larger number of targets in a discriminatory manner without harming uninfected bystander cells. Figure was generated using Biorender.
